# Microbiota-dependent IFN-L controls uterine and placental immunity

**DOI:** 10.64898/2026.01.27.701820

**Authors:** Elisha Segrist, Anne Roman, Eduard Ansaldo, Juliana Perez Chaparro, Colton McNinch, Irini Sereti, Yasmine Belkaid

## Abstract

Precise regulation of uterine immunity is required to support fundamental processes including reproduction and pathogen protection. How the local milieu and constitutive stressors, including the cervicovaginal microbiota, shape the delicate balance underlying uterine immunity is poorly understood. Here, we found that the cervicovaginal microbiota promotes both local immunity and the immunoregulatory activity of interferon lambda (IFN-L) in the uterus. Using murine models, we found a keystone role for IFN-L in constraining the immune tone of this site, in particular of innate lymphoid cells and Th17 cells. Further, in the context of pregnancy, IFN-L enhanced antibacterial responses at maternal-fetal barriers to *Streptococcus agalactiae* infection, thereby controlling fetal and neonatal transmission. Collectively, this work uncovered how IFN-L integrates microbial signals under both steady state and pregnancy conditions and mediates the essential functions of the uterine immune system – antimicrobial protection and immunoregulation.

## Main text

The immune system optimally tunes tissue homeostasis through recognition of and response to the endogenous and exogenous environment. Such regulation is of particular importance at barrier sites like the female reproductive tract (FRT), where the immune system must balance the dynamic and sometimes conflicting requirements for protection against pathogens and fetal tolerance. For instance, FRT immunity is altered by fluctuating levels of sex hormones throughout the menstrual cycle in humans and estrous cycle in mice, a phenomenon that influences vital processes including antimicrobial responses or tissue remodeling and repair^1^. Additionally, local FRT and systemic immunity are profoundly altered throughout gestation^2^. Inflammatory immune processes are required for blastocyst implantation to initiate pregnancy and for the onset of labor to end pregnancy^2, 3^. However, throughout pregnancy, an anti-inflammatory environment characterized by an expansion of regulatory T cells and suppression of the effector activity of T cells is maintained to allow for fetal tolerance and growth^2, 3^.

Work from us and others revealed that all barrier sites are extensively controlled by local tissue factors, including the trillions of bacteria that make up the microbiota^4^. For example, the cervicovaginal microbiota (CVM) has profound effects on women’s health ranging from risk of infection to poor reproductive outcomes^5-7^. The interaction between vaginal microbes and epithelial transcriptional responses has been probed in depth *in vitro*^*8-12*^, but the mechanisms by which the immune system, particularly the uterine immune system, detects the CVM and the downstream processes that this recognition leads to are largely uncharacterized. The existence of an intrauterine microbiota is debated^13-15^, but CVM derived products can shape birth outcomes^16, 17^ suggesting their potential to regulate uterine immunity.

Evolutionarily, protection of the FRT is critically important in maintaining the reproductive capacity of the organism. During pregnancy, sufficient antimicrobial protection is vital to ensure that the substantial maternal investment in supporting fetal development results in healthy offspring. If maternal immunity is either insufficient or too inflammatory then maternal and fetal death or poor birth outcomes can occur^18^. In addition, the inability of the immune system to clear pathogens can result in *in utero*, intrapartum, or perinatal transmission which can cause congenital diseases in offspring^18,19^.

Type III interferons, which in mice include interferon lambda 2 (IFN-L-2) and interferon lambda 3 (IFN-L-3), are uniquely active at barrier sites^20-22^. IFN-L can regulate the microbiota, influence tissue immunity, and protect against viral infection^23-30^. In the gut, homeostatic IFN-L activity is controlled by endogenous stimulation by the microbiota and virome^31, 32^. In the FRT, IFN-L is antiviral in the placenta during pregnancy without inducing pathogenic inflammation leading to fetal death, like type I IFNs^28, 33-41^. Bacteria can induce IFN-L expression, including in the placenta^42^, but the role of IFN-L on placental bacterial control is unknown. Some bacteria, like *Streptococcus agalactiae*, can cause ascending infection of maternal-fetal membranes (chorioamnionitis) which can result in preterm or stillbirth^43-45^. Chorioamnionitis occurs in 1-10% of term births and ∼40% of preterm births and rates of chorioamnionitis are increasing^46-48^. Additionally, offspring are at risk of *in utero* or intrapartum infection which can lead to sepsis, meningitis, or pneumonia^47, 49^. Antibiotics and expedited delivery are performed after chorioamnionitis is detected, but exposed infants can experience long-term health consequences^50^. While transmission and infection risk are multifactorial, the maternal and neonatal immune parameters that shape transmission events remain largely unknown^51^.

Here, we show that, consistent with the need to tightly regulate uterine immunity, the CVM promoted both local immunity and immune regulation within the uterus. Indeed, IFN-L constrained type 17 uterine immune abundance in a microbiota-dependent manner. Further, IFN-L played an important antibacterial role during pregnancy which influenced *in utero* transmission and intrapartum infection of offspring. Collectively, these data uncovered a new role for the CVM and IFN-L in the control of a fundamental reproductive organ.

### FRT colonization by commensal bacteria promotes type 1 and type 17 immune responses in the uterus

Sex hormone levels fluctuate throughout the murine estrous cycle, with a peak of estrogen during proestrus and a peak of progesterone during diestrus (**Fig 1A**)^52^. Since estrogen and progesterone can alter local immunity in distinct ways^1, 52, 53^, we treated animals with either 17β-estradiol (estrogen) or medroxyprogesterone acetate (progesterone) to stabilize the uterine environment throughout the course of an experiment by maintaining animals in either the proestrus or diestrus stage of the estrous cycle.

**Figure 1.**
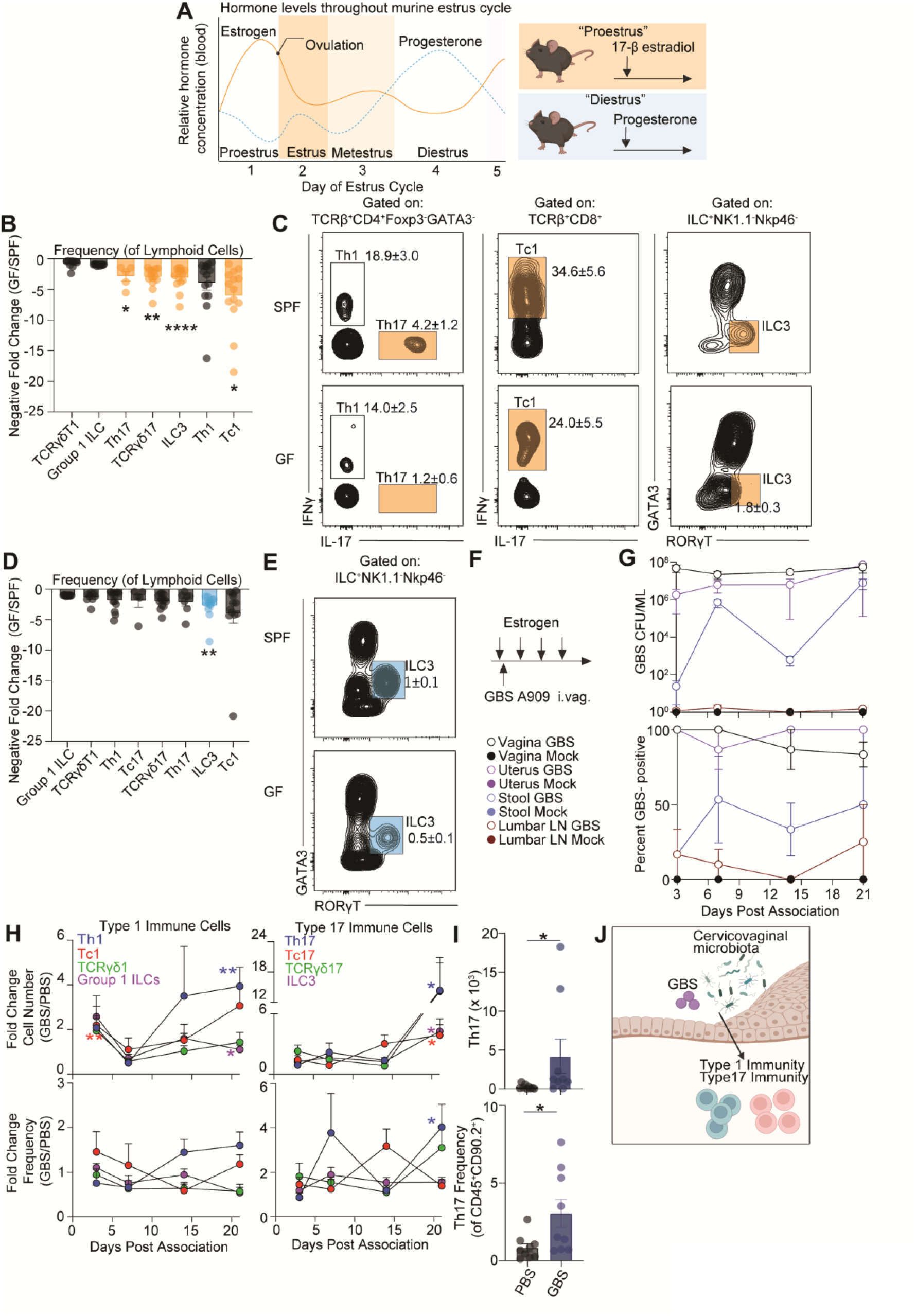
The local microbiota regulates type 1 and type 17 immunity in the uterus. (A) Schematic of the mouse estrus cycle adapted from a BioRender.com template and Walmer *et al*. (B) Quantification of type 1 and type 17 immune cells shown as negative fold change cell frequency (of CD45^+^CD90.2^+^ cells) in GF animals compared to SPF animals treated with estrogen. (C) Representative FACS plots displaying frequency of uterine Th1, Th17, Tc1, Tc17, and ILC3 cells in estrogen treated SPF and GF animals. (D) Quantification of type 1 and type 17 immune cells shown as negative fold change of cell frequency (of CD45^+^CD90.2^+^ cells) in GF animals compared to SPF animals during progesterone treatment. (E) Representative FACS plots displaying frequency of uterine ILC3 cells in SPF and GF animals following progesterone treatment. (F) Experimental schematic showing intravaginal commensal association of GBS A909 after estrogen treatment of nulliparous mice. (G) Top, quantification of GBS bacterial burden in the vagina, uterus, stool, and lumbar lymph nodes (LN) over the 21 dpa time course. Bottom, GBS colonization rate in the vagina, uterus, stool, and lumbar lymph nodes over the 21 dpa time course. (H) Left, quantification of uterine type 1 immune cells shown as fold change of absolute cell number and cell frequency (of CD45^+^CD90.2^+^ cells) in GBS associated animals compared to mock treated animals. Right, quantification of uterine type 17 immune cells shown as fold change in absolute cell number and cell frequency (of CD45^+^CD90.2^+^ cells) in GBS associated animals compared to mock treated animals. (I) Quantification of Th17 absolute cell number and cell frequency (of CD45^+^CD90.2^+^ cells). (J) Working model. (B, D, I) Each dot represents an individual tissue with mean ± SEM from at least (I) two or (B, D) three biological replicates. (H) Mean with ± SEM from at least two biological replicates. (B, D, H, I) Significance was determined by an unpaired Student’s t-test. * denote p-val<0.05 ** denote p-val<0.01 **** denote p-val<0.0001.

We used flow cytometry to characterize immune cell abundance and activation in the uterine compartment of progesterone or estrogen treated mice (**Fig S1A, S1B**). The frequency and number of lymphoid cells was higher in the uteri of progesterone treated mice compared with estrogen treated mice (**Fig S2A, S2B**). Lymphoid cells represented ∼36% or ∼21% of hematopoietic cells in progesterone or estrogen treated mice, respectively (**Fig S2B**). As previously shown^54^, the tone of the uterine compartment throughout the estrous cycle was skewed toward type 1 immunity with Tbet^+^ group 1 innate lymphoid cells (ILCs; primarily composed of tissue resident NK cells) representing the majority (∼45-59%) of the lymphoid compartment (**Fig S2C, S2D**). The abundance of uterine Group 1 ILCs was higher in progesterone treated mice compared to estrogen treated mice (**Fig S2D**). Type 17 immune cells (RORγT^+^ or IL-17^+^ cells) were the next most abundant cell type, comprising about ∼8-11% of the lymphoid compartment (**Fig S2C**). The uterine frequency of IL-17^+^ γδ T cells (TCRγδ17) was higher in progesterone treated animals relative to estrogen treated animals, whereas the frequency of ILC3, Th17 and Tc17 cells was unchanged between the conditions (**Fig S2E**). Type 2 immune cells (GATA3^+^) were the least abundant lymphoid cell type (**Fig S2C**) and the abundance of uterine ILC2 and Th2 cells was the same between estrogen and progesterone treated animals (**Fig S2F**). We next focused on uterine type 1 and type 17 immune programs as they are highly abundant in the uterus and are significantly influenced by the microbiota at other body sites^4, 55-60^.

To understand how the microbiota could influence uterine immunity, we first compared the number and activity of lymphoid and myeloid cells in specific pathogen free (SPF) or germ-free (GF) mice using flow cytometry. During estrogen treatment, we found that SPF mice exhibited higher frequencies of uterine Th17, TCRγδ17, and ILC3 cells relative to GF animals (**Fig 1B, 1C)**. Additionally, there was a higher number of uterine Th17 cells in SPF mice relative to GF mice (**Fig S3A**). While SPF mice had a reduced frequency of Tc1 cells and a trend toward reduced Th1 cells relative to GF mice, there was no effect on the frequency of Group 1 ILCs or IFNγ^+^ γδ T cells (TCRγδ1) (**Fig 1B**). When we examined the effect of the microbiota on the myeloid compartment, we found no significant differences in absolute cell number or frequency of macrophages, monocytes, type 1 or type 2 dendritic cells (DC1, DC2) or neutrophils between SPF and GF uteri (**Fig S3B**).

In progesterone treated mice, the impact of the microbiota on the lymphoid compartment was less pronounced with only a significant reduction of ILC3 frequency in GF mice (**Fig 1D, 1E, S3C**). GF mice also had increased frequencies of neutrophils relative to SPF mice (**Fig S3D, S3E**) suggesting a potential role for the microbiota in the control of the inflammatory tone of the tissue within this specific hormonal state.

As GF mice lack a microbiota at all barrier sites, these results did not distinguish whether the observed effect was due to local or systemic action of the microbiota. The CVM and its associated products can alter reproductive outcomes, suggesting the ability to influence the uterine environment^5, 10, 16, 17^. To test the possibility that the CVM could directly control uterine immunity, we colonized SPF mice with the vaginal commensal, *S. agalactiae* (GBS), commonly found in the human CVM^61, 62^ (**Fig 1F**). We used GBS as a model as it readily colonizes the murine FRT^63^ unlike other gram-positive commensals^64-67^ and has been used extensively to identify bacterial factors that shape colonization and ascending infection^68, 69^. Mice in diestrus are not stably colonized with GBS, so continuous treatment with estrogen is required to establish and maintain GBS colonization in the murine vagina^63, 70^. By 3 days post-association (dpa), GBS strain A909 robustly colonized the vagina and uterus with up to 10^8^ CFU/mL found in nearly all animals and colonization was maintained throughout the 21 day experiment (**Fig 1G**). Since GBS is also found in the gastrointestinal tract^71, 72^, we monitored the stool for GBS and found varying levels of GBS burden in about 50% of all animals (**Fig 1G**). Intermittent and minute levels of bacteria were found in the lumbar lymph nodes that drain from the uterus confirming that this model of association did not cause infection (**Fig 1G**).

Evaluation of lymphoid changes between mock and GBS associated animals revealed alterations in the activation and frequency of subsets of ILCs, γδ T cells, and αβ T cells at 21 dpa. Specifically, exposure to a new commensal within the FRT (vagina and uterus) resulted in a significant increase in type 17 immune cell numbers (Th17, Tc17, ILC3) when represented as fold change relative to mock treated animals at 21 dpa (**Fig 1H, 1I, S3F**). There was also an increase in type 1 immune cells by 21 dpa, with a significant fold increase in Th1 cell number (**Fig 1H, S3F**).

We next assessed if the impact of GBS association on type 1 and type 17 immune cells was mediated by live bacteria as opposed to bacterially derived products. To mimic the persistent antigen exposure associated with robust GBS colonization, mice were treated daily with 10^8^ CFU heat killed GBS (HK GBS) or GBS spent media (1x or 10X) (**Fig S4A**). A single administration of PBS, GBS, or daily administration of Todd Hewitt bacterial growth media (THB) were also included as controls. GBS supernatant promoted IL-17 production from CD4^+^ T helper cells similarly to live GBS (**Fig S4B, S4D, S4E**) which supported the idea that microbial derived products are sufficient to promote local immune responses. On the other hand, only live bacteria promoted IFNγ production from CD4^+^ T helper cells (**Fig S4C, S4D, S4F**). Thus, bacterial products were sufficient to promote uterine type 17 immunity, while type 1 immunity was only induced by live CVM.

By using two complementary murine models to probe the effects of the microbiota on the uterine immune environment, we found that the CVM promoted type 1 and type 17 immune cell accumulation and cytokine expression (**Fig 1J**).

### IFN-L constrains uterine immune abundance in a microbiota-dependent manner

We and others have previously shown that the microbiota promoted type I IFN signaling that in turn influenced T cell responses^57, 73-77^. Therefore, we investigated whether type I IFNs alter homeostatic uterine type 17 immunity. To this end, we temporally depleted type I IFN signaling by treating SPF female mice with a blocking antibody against the type I IFN receptor (IFNAR) (**Fig S5A**). Temporal blockade of type I IFN signaling had no effect on the abundance or function of type 17 lymphocytes (**Fig S5B**). These data suggest that a different innate immune pathway bridges microbiota detection and uterine immune regulation.

As IFN-Ls are induced by the microbiota and are immunoregulatory^20, 22, 31, 32^, we next considered a potential role for IFN-L in regulating uterine immunity. Since estrogens dampen IFN signaling^78-80^, we focused on the effect of IFN-L in progesterone treated animals. IFN-L signaling was temporally blocked using a neutralizing antibody against IFN-L-2/3 (a-IFN-L)^79, 81, 82^ in SPF or GF adult female mice (**Fig 2A**). Loss of IFN-L signaling in SPF animals resulted in a significant increase in the abundance of type 17 immune cells, including Th17, RORγT^+^ γδ T cells, and ILC3s relative to isotype control (Iso) treated animals (**Fig 2B, 2C, 2D, S6A**). Furthermore, we found that IFN-L’s regulation of immunity is dependent on the microbiota, as loss of IFN-L signaling in GF mice had no effect on the abundance of Th17, RORγT^+^ γδ T cells, and ILC3s (**Fig 2C, 2D, S6A**). Thus, these data show that IFN-L constrains type 17 uterine immunity in a microbiota-dependent manner.

**Figure 2.**
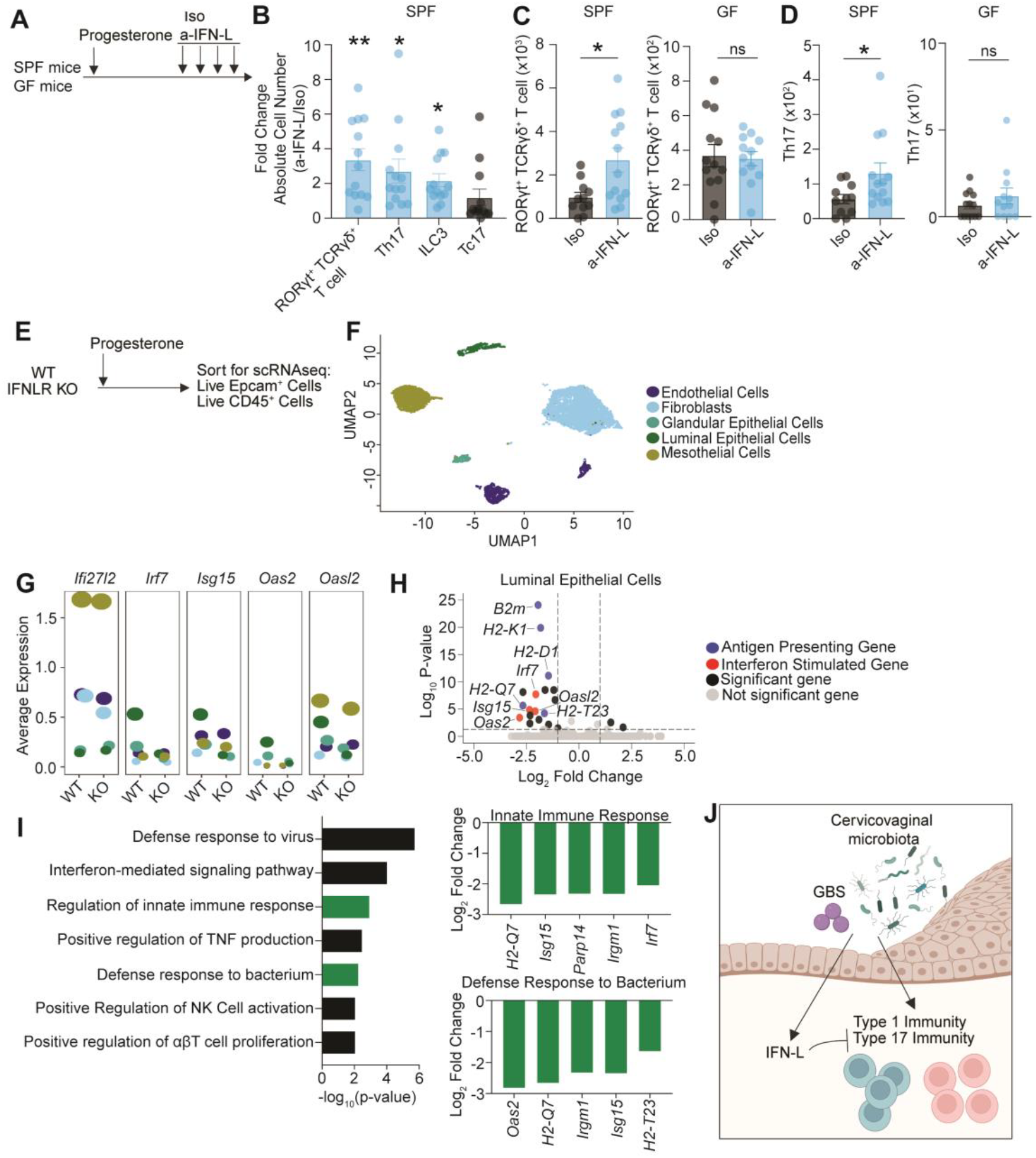
IFN-Ls constrain uterine homeostatic immunity in a microbiota-dependent manner. (A) Experimental schematic depicting isotype control (Iso) and a-IFN-L blocking antibody treatment (a-IFN-L) following progesterone administration in SPF or GF mice. (B) Quantification of uterine type 17 lymphoid cells shown as fold change in absolute cell number in progesterone treated SPF animals treated with a-IFN-L relative to Iso treated animals. (C, D) Quantification of absolute cell number of uterine (C) RORγT^+^ γδT cells and (D) Th17 cells. (E) Experimental schematic showing progesterone treatment of wild type (WT) littermate control or IFN-L receptor (IFNLR) knock out mice, followed by sorting of EpCam^+^ and CD45^+^ cells for single cell RNA sequencing. (F) UMAP projection showing uterine cells sorted using an EpCam gate. (G) Expression of the ISGs *Ifi27l2a, Irf7, Isg15, Oas2, Oasl2* across EpCam^+^ clusters from WT and IFNLR KO mice. (H) Volcano plot displaying differentially expressed genes from uterine luminal epithelial cells. (I) Top enriched GO term pathways for genes downregulated in uterine luminal epithelial cells from IFNLR KO mice, with Log_2_ fold change values shown for differentially expressed genes in each highlighted pathway. (J) Working model. (B-D) Each dot represents an individual tissue from three biological replicates with mean with ± SEM. Significance determined by an unpaired Student’s t-test. * denote p-val <0.05 ** denote p-val <0.01.

### IFN-L controls interferon stimulated gene (ISG) expression in luminal epithelial cells and fibroblasts

To identify IFN-L responsive cells and the potential homeostatic function of IFN-L in the uterus, we performed single cell RNA sequencing of sorted EpCam^+^ and CD45^+^ cells from littermate control wild type (WT) and IFNLR KO adult female mice (**Fig 2E, 2F, S6B, S6D, S6E**). As expected, IFNLR expression (*Ifnlr1*) was limited compared with the ubiquitous type I IFN receptor (*Ifnar1*) (**Fig S6C**)^83^. Notably, IFNLR was primarily expressed by uterine luminal and glandular epithelial cells (**Fig S6C**). IFNLR expression was also detected in DCs and macrophages but was absent from other CD45^+^ cells.

We observed reduced ISG expression only in uterine luminal epithelial cells (*Irf7, Isg15, Oas2*, and *Oasl2*) and in fibroblasts (*Ifi27l2a*) from IFNLR KO mice relative to WT mice (**Fig 2G, 2H**). These data suggest that while several cell types express IFNLR, only luminal epithelial cells and fibroblasts tonically respond to IFN-Ls (**Fig S6C, 2G, 2H**). In the absence of IFNLR, luminal epithelial cells also had reduced expression of genes related to antigen presentation (**Fig 2H**), a common feature of IFN signaling^84^. GO term analysis of genes downregulated in uterine luminal epithelial cells from IFNLR KO mice revealed that IFN-L signaling promoted expression of genes involved in pathways like innate immune responses, defense responses to bacteria and viruses, regulation of natural killer cell activation, and regulation of T cell proliferation (**Fig 2I**). These data indicate that luminal epithelial cells express high levels of IFNLR and that loss of homeostatic IFNL signaling results in reduced expression of ISGs and genes involved in antimicrobial defense.

These data suggest that the microbiota both promotes local uterine immunity and controls IFN-L immunoregulator activity, which constrains uterine type 17 immunity (**Fig 2J**). Furthermore, our scRNAseq data suggest that IFN-L may be involved in antibacterial defense responses.

### IFN-L has no impact on bacterial burden in non-pregnant animals

Next, we wanted to probe IFN-L’s function in the uterus. While IFNs are primarily investigated in the context of viral infection, growing evidence implicates IFNs in immune responses to bacteria as well^22, 85^. For instance, IFN-L regulates the nasal microbiota community structure and can either promote lung infection with pathogens like *S. aureus* and *B. pertussis* or protect against bacterially induced pneumonia^23, 86-88^. The role of IFN-L in bacterial defense within the FRT is unknown. Since IFN-L controls expression of genes related to bacterial defense in uterine luminal epithelial cells (**Fig 2I**), we next investigated IFN-L’s role in controlling bacterial burden of the vaginal commensal GBS following intravaginal association in nulliparous animals (**Fig 3A**). There was no difference in GBS burden in the vaginal lavage of animals treated with a-IFN-L or Iso blocking antibody at any time point after vaginal colonization (**Fig 3B**). Additionally, there was no difference in GBS burden in vaginal or uterine tissues from a-IFN-L or Iso treated mice at 6 dpa (**Fig 3C, 3D**). These findings indicate that IFN-L did not regulate bacterial abundance in the uterus and vagina. These results agreed with previous studies showing that basal IFN-L signaling was not antiviral against Zika virus in the nulliparous WT murine uterus^33, 34^ and did not protect against herpes simplex virus or murine papillomavirus infection of the vagina^83, 89^. Overall, these data indicate that, unlike in the gut and the lungs, homeostatic IFN-L is not involved in microbial defense in the nulliparous uterus^22-25, 27, 29-34, 83, 86-89^.

**Figure 3.**
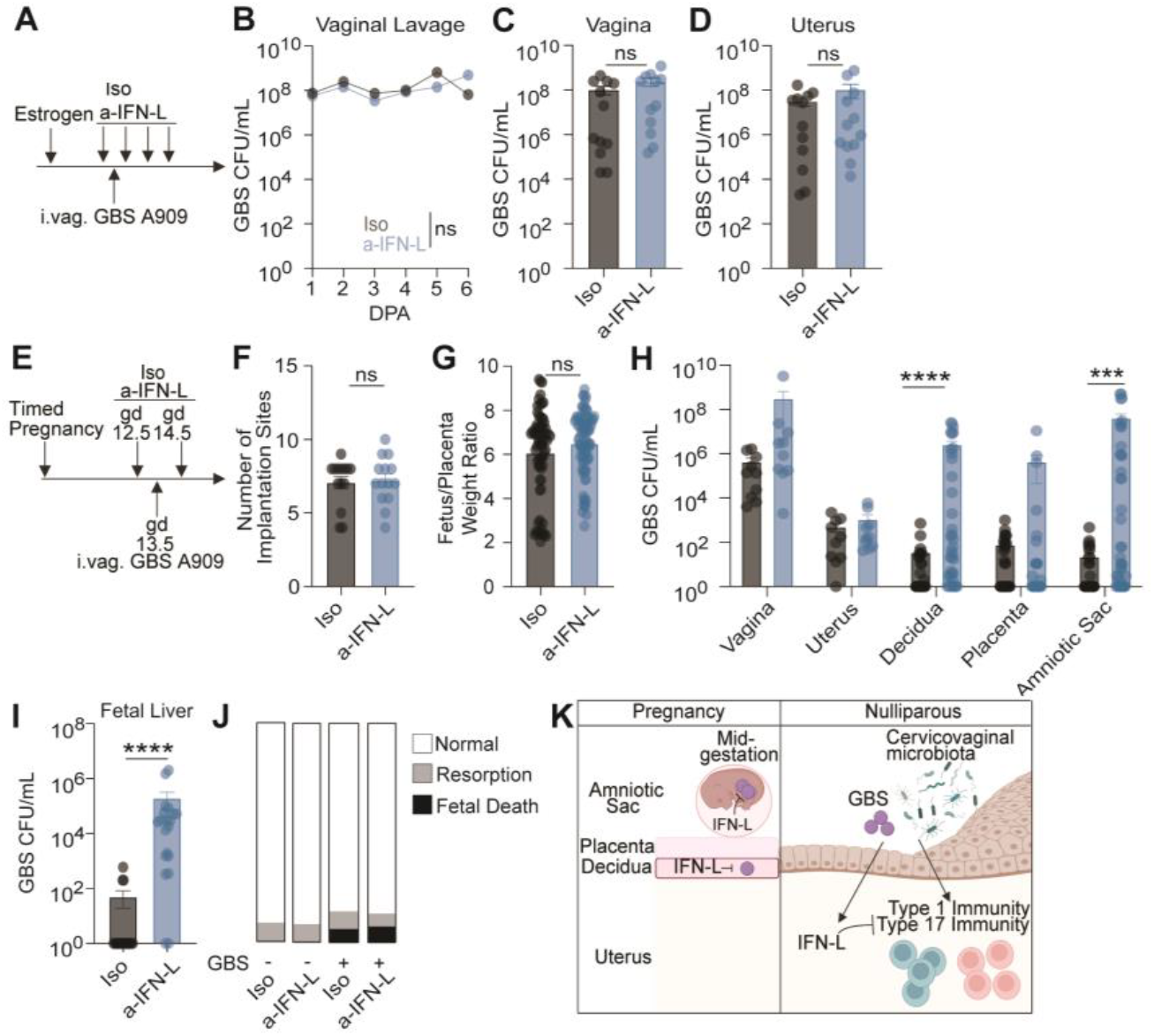
IFN-L is antibacterial specifically during pregnancy. (A) Experimental schematic showing Iso or a-IFN-L treatment before and throughout intravaginal GBS colonization of nulliparous animals. (B-D) Quantification of GBS counts in (B) vaginal lavages from 1-6 dpa, (C) vaginal and (D) uterine tissue at 6 dpa. (E) Experimental schematic showing Iso or a-IFN-L treatment before and throughout intravaginal GBS infection of timed pregnant dams. (F) Number of implantation sites per litter in Iso or a-IFN-L treated GBS-infected pregnant dams. (G) Quantification of the fetus-to-placenta weight ratio in Iso or a-IFN-L treated GBS-infected pregnant dams at gd16.5. (H) Quantification of GBS counts in indicated tissues at 3 dpi. (I) Quantification of GBS counts in fetal liver at 3 dpi. (J) Graphical representation of fetal outcomes found in mock or GBS-infected Iso or a-IFN-L treated pregnant dams at gd16.5. (K) Working model. (B) Significance was determined by two-way ANOVA with Tukey’s multiple comparison test with mean ± SEM from three biological replicates. (C, D, G, H, I) Each dot represents an individual tissue from at least three biological replicates with mean ± SEM. (F) Each dot represents the total number of implantation sites per individual dam from at least three biological replicates with mean ± SEM. (C, D, F-I) Significance determined by an unpaired Student’s t-test. *** denote p-val<0.001, **** denote p-val<0.0001.

### IFN-L controls GBS infection and *in utero* transmission during pregnancy

Previous studies revealed that IFN-L is antiviral during mid to late pregnancy in both mice and humans^33, 36, 39^. While bacterial infection induced IFN-L expression in the murine placenta^42^, it is unknown if IFN-L controls bacterial infection of maternal-fetal barriers. GBS is present in the CVM of ∼30% of women with no negative health effects^61, 62^. However, during pregnancy this bacterium can become pathogenic, causing ascending infection of maternal-fetal membranes^43, 45, 47^. To define the role of IFN-L during bacterial infection in pregnancy, we used an ascending model of GBS infection during mid-gestation that results in chorioamnionitis^90, 91^. Timed pregnancies were established to ensure synchronized gestations and mice were treated with an a-IFN-L or Iso blocking antibody before and throughout GBS infection (**Fig 3E**). Treatment with a-IFN-L during GBS infection had no effect on litter size or the fetal-to-placenta weight ratio (**Fig 3F, 3G**). At 3 days post infection (dpi), GBS burden was quantified in vaginal swabs and uterine, decidual, placental, and amniotic tissues. We found that neutralization of IFN-L signaling resulted in increased GBS burden in the decidua and amniotic sac with no effect on GBS burden in the vagina, uterus or placenta (**Fig 3H**). While GBS infected the maternal-fetal barriers of Iso treated control dams, *in utero* transmission was rare (**Fig 3I**). In contrast, fetuses of dams treated with a-IFN-L during GBS infection had significantly higher levels of GBS in the fetal liver (**Fig 3I**). Despite the presence of *in utero* transmission in dams treated with a-IFN-L during GBS infection, we observed no increase in early resorption or fetal death (**Fig 3J**). In all, these data indicate that during pregnancy, IFN-L protected the decidua and fetus from ascending bacterial infection (**Fig 3K**).

### IFN-L-dependent antibacterial activity *in utero* influences offspring infection

We found that loss of IFN-L activity during bacterial infection enhanced *in utero* GBS transmission without increasing fetal demise (**Fig 3I, 3J**), so we next investigated the effects on birth outcomes and neonatal health. To do this, we established timed pregnancies, treated dams with Iso or a-IFN-L blocking antibody throughout pregnancy and infected dams with GBS (**Fig 4A**). In these experiments, we allowed dams to give birth and monitored maternal survival, GBS carrier status and neonatal weight and infection. There was minimal maternal mortality before or after delivery in Iso or a-IFN-L treated GBS infected dams and all dams cleared GBS from the vagina by postnatal day 11 (**Fig 4B, 4C**). GBS infection and treatment with Iso or a-IFN-L had no significant effect on litter size, consistent with the lack of fetal death caused by GBS infection (**Fig 4D, 3J**). We monitored neonatal weight after birth and found no significant differences among offspring born to mock infected dams, GBS infected dams or GBS infected dams with deficient IFN-L signaling (**Fig 4E**).

**Figure 4.**
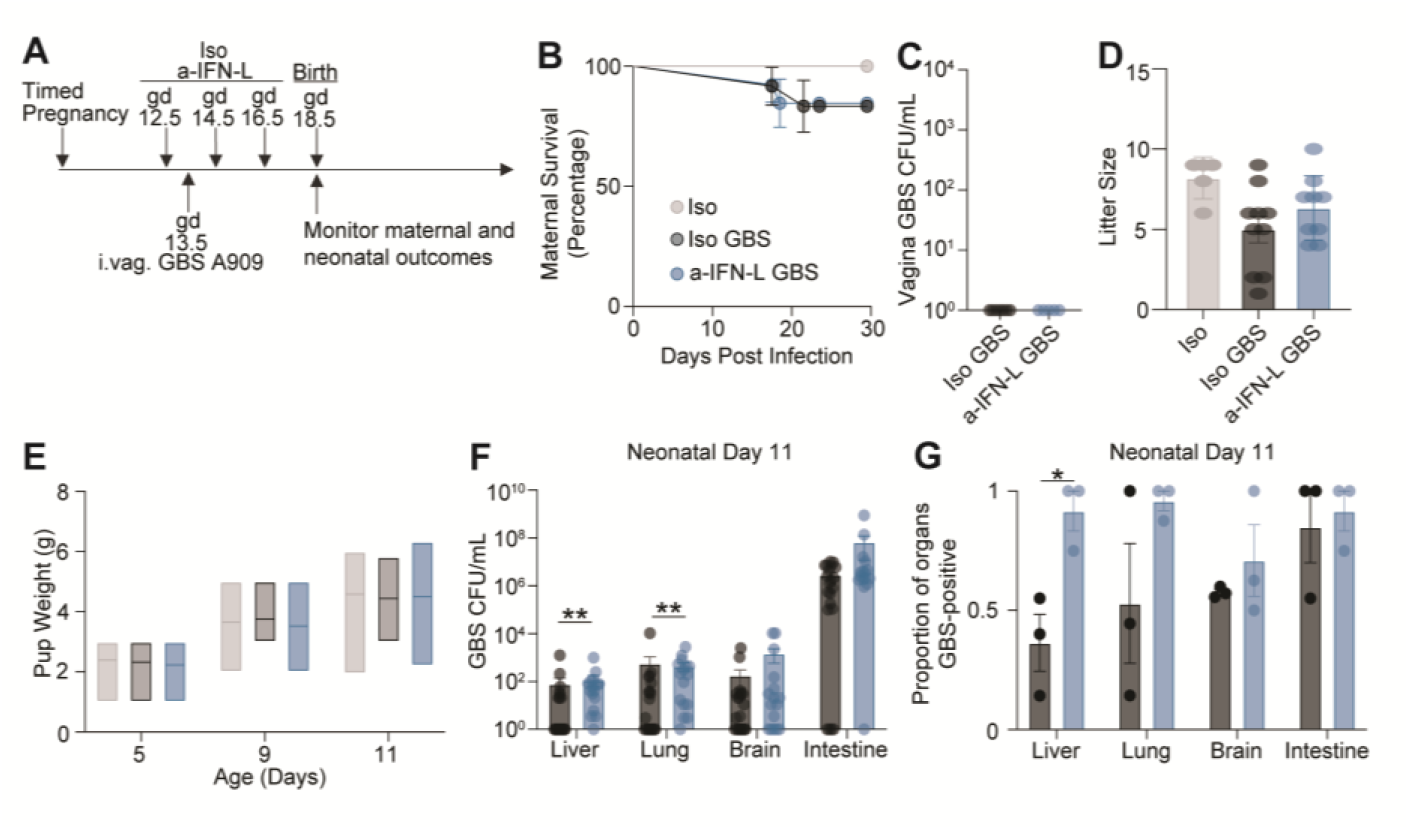
Neonates born from GBS infected dams deficient in IFN-L signaling have enhanced GBS infection. (A) Experimental schematic showing Iso or a-IFN-L neutralizing antibody treatment before and throughout intravaginal GBS infection of timed pregnant dams. Maternal and neonatal outcomes were monitored through perinatal day 11. (B) Survival curve displaying maternal survival following GBS infection during pregnancy and through the first 11 days of the perinatal period. (C) Quantification of vaginal GBS burden in dams at perinatal day 11. (D) Quantification of litter size per individual dam at postnatal day 5. (E) Quantification of neonatal weight at postnatal day 5, 9 and 11. (F) Quantification of GBS CFU/mL in the neonatal liver, lung, brain, and intestine at postnatal day 11. Each dot represents an individual tissue with mean ± SEM. (G) Proportion of neonatal livers, lungs, brains, and intestines positive for GBS at postnatal day 11. (B-G) Three independent replicates were performed. (B) A log-rank test was performed to determine statistical significance. (C, F, G) An unpaired Student’s t-test was performed to determine statistical significance. (D) A one-way ANOVA with multiple comparisons was performed to determine statistical significance. (E) A two-way ANOVA with multiple comparisons was performed to determine statistical significance. *denote p-val <0.05.

We assayed body sites in neonatal offspring that have been previously shown to be infected during the intrapartum period by GBS^92^. Neonates born to GBS infected dams treated with Iso or a-IFN-L had similar colonization rates and GBS burdens in the gastrointestinal tract, likely reflecting colonization that occurs at birth^93^. In contrast, neonates born to GBS infected a-IFN-L treated dams had higher infection levels in the lungs and liver compared with neonates born to GBS infected Iso treated dams (**Fig 4F**). Furthermore, there was a higher proportion of GBS-positive livers in neonates born from GBS infected a-IFN-L treated dams compared with neonates born from GBS infected Iso treated dams (**Fig 4G**). These data are consistent with our observation that loss of IFN-L during GBS infection in pregnancy resulted in increased *in utero* transmission to fetuses (**Fig 3I**). Together, these results indicate that loss of IFN-L signaling throughout GBS infection in pregnancy enhanced GBS infection of the lungs and livers in offspring.

## Discussion

Here, we described a role for the CVM in mediating uterine immune homeostasis by promoting local uterine immunity while simultaneously engaging IFN-L-mediated immunoregulatory mechanisms. We also identified a previously unknown role for IFN-L in antibacterial defense in pregnancy which shaped *in utero* transmission and intrapartum infection of offspring.

Recent findings in humans have revealed that CVMs dominated by Gram-negative commensals have increased activation of cervical type 17 immune cells compared to women with low diversity CVMs dominated by Gram-positive commensals^94, 95^. We find that the cervicovaginal Gram-positive commensal GBS which is found in women with either low or high diversity CVMs^62^, induces type 17 and type 1 immune accumulation in the uterus (**Fig 1B-E, 1H, 1I**). Furthermore, we find that mice without any microbiota have reduced type 17 and type 1 uterine immune responses compared to mice with an endogenous microbiota. These data suggest that cervicovaginal bacteria promote type 17 immune responses, while specific commensals or CVM communities may drive excessive or dysregulated type 17 immune activation in the FRT.

Dysregulation of type 17 immunity is associated with endometriosis^96-98^ and pregnancy complications^99-102^ underscoring the necessity of precise regulation of this pathway in the FRT. Endometriosis and pregnancy complications, like pre-term birth, are linked to vaginal dysbiosis or to specific vaginal commensals, like GBS^5, 10, 43, 103-105^. We find that while the microbiota promotes type 17 immune accumulation in the uterus, it also engages a second IFN-L-dependent mechanism that constrains type 17 immune responses (**Fig 2B-D, S6A**). Therefore, the connection between the microbiota and pathogenic type 17 immunity in these pathologies should be further explored.

As loss of IFN-L resulted in enhanced type 17 immune responses in a microbiota-dependent manner, we propose that IFN-L is a regulator of type 17 immune responses. We recognize that an alternative hypothesis is that loss of IFN-L may cause an outgrowth of bacteria which then induce uterine type 17 immune responses. However, when we evaluated whether IFN-L influences GBS burden in nulliparous mice, we found that mice deficient in IFN-L signaling had similar GBS bacterial burden relative to mice with intact IFN-L signaling (**Fig 3A-D**). Thus, our data suggest that IFN-L regulates uterine type 17 immune responses independently of CVM bacterial burden, consistent with a model in which IFN-L acts on host pathways rather than indirectly influencing immunity through CVM alterations.

The incidence of ascending GBS infection in pregnancy is low despite a relatively high GBS carrier rate in pregnant women^43, 45, 47, 49, 61, 72, 106^. While the basis of this discrepancy is likely multifactorial, our data revealed a previously unrecognized role for IFN-L in protecting against chorioamnionitis and in utero GBS transmission (**Fig 3H, 3I**). Whether IFN-L’s antibacterial activity in pregnancy extends to bacterial pathogens like *Listeria monocytogenes* which causes severe infection in pregnancy through a hematogenous route of infection remains to be determined^107^.

Additionally, genetic polymorphisms in human IFN-L genes are associated with HCV clearance and Zika virus outcomes in pregnancy^40, 108^. Given our findings, it will be important to determine whether IFN-L genetic polymorphisms influence susceptibility to ascending GBS infection during pregnancy.

Our data indicate that IFN-L’s antibacterial action *in utero* influences GBS infection of neonates (**Fig 4F, 4G**). Children born to GBS positive women are more likely to develop asthma which has been attributed to prophylactic antibiotic treatment of the mother during delivery^50^. We find that neonates born from dams deficient in IFN-L signaling during GBS infection in pregnancy have increased GBS burden in organs including the lungs relative to neonates born from dams with no defect in IFN-L signaling during GBS infection (**Fig 4F, 4G**). These data raise the possibility that IFN-L influences GBS transmission and neonatal lung health even in the absence of early life antibiotic exposure.

Overall, our findings reveal that IFN-L orchestrates uterine homeostasis and extends the known antiviral role of IFN-L in pregnancy to include protection against bacterial infection. This work provides new insight into the optimal control of uterine immunity, highlighting the necessity of balancing microbiota-dependent cellular immune accumulation and IFN-L-mediated immune regulation.

## Materials and Methods

### Mice

Conventional SPF C57BL/6J WT mice were bred, maintained, and obtained by the NIAID Taconic Exchange Program. Germ free C57Bl/6 mice were bred and maintained in the National Allergy and Infectious Diseases (NIAID) Microbiome Program gnotobiotic animal facility and in the National Cancer Institute’s Frederick National Laboratory gnotobiotic animal facility. Ifnlr^-/-^ mice were obtained from the Sergei Kotenko Laboratory^109^ and were bred and maintained under SPF conditions. Mice were provided a standard pelleted rodent diet; in experiments with SPF and GF mice, SPF mice were maintained on the standard pelleted rodent diet used in the gnotobiotic facility. All mice were bred and maintained at an American Association for the Accreditation of Laboratory Animal Care (AAALAC)–accredited animal facility at NIAID and housed in accordance with the procedures outlined in the Guide for the Care and Use of Laboratory Animals. Age-matched SPF mice between 8-12 weeks of age were used for each experiment. All experiments were performed at NIAID under an Animal Study Proposal (LHIM2E) approved by the NIAID Animal Care and Use Committee.

### Streptococcus agalactiae culture

*S. agalactiae* A909 (ATCC BAA-1138) was grown for 16 hours on selective CHROMagar StepB (DRG International, Inc.) agar plates. A909 was then subcultured in Todd-Hewitt broth (NutriSelect® Plus) and incubated at 37^°^C until mid-logarithmic phase (OD_600_ = 0.4-0.6). Bacterial cultures were centrifuged at 5000 rpm for 5 minutes, washed in sterile PBS, and resuspended in sterile PBS at 10^9^ CFU/mL (nulliparous mice) or 10^10^ CFU/mL (pregnant mice).

### Sex hormone treatment

Mice were injected subcutaneously in the neck scruff with 100 μL of a 20 mg/mL solution of medroxyprogestrone acetate (Prasco Laboratories) in PBS and referred to as progesterone treated throughout the manuscript. Mice were injected subcutaneously in the neck scruff with 100 μL of a 1 ng/μL solution of beta-estradiol (Sigma Aldrich #E2758) in corn oil and referred to as estrogen treated throughout the manuscript. Mice were maintained on estrogen treatment every four days for the length of the experiment. For *S. agalactiae* association experiments, nulliparous mice were treated with 100 μL of a 5 mg/mL solution of beta-estradiol a day before bacterial administration as previously described^63^. *S. agalactiae* associated mice were maintained on estrogen treatment every seven days for the length of the experiment.

### S. agalactiae intravaginal association in nulliparous mice

One day after estrogen treatment, animals were intravaginally administered 10^9^ CFU/mL of *S. agalactiae* A909 in 10 μL of sterile PBS using a gel-loading pipette^63^. Mice were walked for approximately 60 seconds to prevent spilling of the inoculum.

To determine colonization rates and *S. agalactiae* abundance after intravaginal inoculation, we collected vaginal swabs or vaginal lavages, uteri, and stool at 3, 7, 14, and 21 dpa. To evaluate the ability of A909 to disseminate from the FRT we also collected the lumbar draining lymph nodes at each time point.

For antibody neutralization experiments, mice were treated with a neutralizing antibody or isotype control antibody one day before *S. agalactiae* association. Treatment with a neutralizing antibody was maintained every other day throughout the experiment. *S. agalactiae* CFU/mL was quantified in the vaginal lavage at 1-6 dpa and in the vaginal and uterine tissue at 6 dpa.

### Timed pregnancies

WT mice were single housed for 1-2 weeks to accumulate dirty bedding and to allow for sperm recuperation. WT females were exposed to dirty bedding from males for 2-3 days to induce estrus. They were then housed with male mice overnight and the presence of a vaginal plug was checked the next morning^53^. If a vaginal plug was found, then mice were considered to be at gestation day (gd) 0.5 and separated from the male mice. Mice found without a plug were allowed to mate another day so the gestation day in all experiments has a range of (±) 1 day of gestation.

### S. agalactiae intravaginal infection of pregnant mice

On gd13.5, mice were intravaginally inoculated with 10^10^ CFU/mL of *S. agalactiae* in 10 μL of sterile PBS^91^. For bacterial quantification experiments, vaginal swabs, uteri, deciduas and metrial glands (n=2-4/dam), placentas (n=2-4/dam) amniotic sacs (n=2-4/dam), and fetal livers (2-4/dam) were collected at 3 dpi. Corresponding organs from mock infected animals were also collected as a negative control.

For antibody neutralization experiments during pregnancy, mice were treated with blocking or isotope control antibodies at gestation day 12.5 and every other day for the course of the experiment.

The impact of infection on fetuses was evaluated at 3 dpi. The presence of fetal death or early resorptions was recorded and quantified. No pre-term births were detected before the 3-day post infection time point.

The impact of infection on neonatal offspring health was evaluated through neonatal day 11. The impact on offspring weight was evaluated at neonatal day 5, 9, and 11. The impact on offspring infection was evaluated at 11 days old. The livers, lungs, brains, and gastrointestinal tracts were collected from 2-4 offspring per dam.

### Quantification of S. agalactiae counts

Quantification of *S. agalactiae* bacterial burden was performed as previously described^63^. A sterile calcium alginate swab (Puritan™) was dipped in sterile PBS and inserted in the vagina of mock infected or GBS associated or infected mice. The swab was turned 5 times clockwise and 5 times counterclockwise with the swab continuously maintaining contact with the walls of the vagina. The swab was then placed in sterile PBS and vortexed at high speeds for 15 seconds to dislodge bacteria from the swab. For vaginal lavages, 100 μL of sterile PBS was added to the vaginal canal and the volume was pipetted up and down 10 times. The nulliparous uterus, vagina, stool and lumbar lymph nodes were homogenized using a bead-based homogenization system (Precellys). Serial tenfold dilutions were performed to quantify bacterial burden using selective CHROMagar StepB (DRG International, Inc.) agar plates. For the lymph nodes, the entire 100 μL sample of undiluted homogenate was plated on CHROMagar Strep B plates.

During and after pregnancy, the vagina was swabbed and *S. agalactiae* was quantified as above. The uterine horns were homogenized using a probe-based tissue homogenizer. All other tissues were homogenized using a bead-based homogenization system (Precellys). Tenfold serial dilutions were plated on CHROMagar Strep B plates. For tissues with low bacterial burden, 100 μL of undiluted homogenate were plated on CHROMagar Strep B plates.

### In vivo treatment with blocking antibodies

Mice were treated with sex hormones and then injected intraperitoneally with 0.5 mg of anti-mouse IFNAR1 antibody (clone MAR1-5A3, BioXcell) or mouse IgG1 isotype control (clone MOPC-21, BioXcell) and then 0.25mg every two days for the duration of the experiment^110^. Mice were also injected intraperitoneally with 2.25 mg/kg of anti-mouse IFN-L-2/3 antibody (MAB17892, R&D Systems) or rat IgG_2b_ isotype control antibody (clone MAB0061, R&D systems) as previously published^79, 81, 82^.

### Murine Tissue Processing for single cell suspensions

Uterine horns were excised and minced into 2 mm pieces and digested in digestion media (RPMI media supplemented with 2 mM L-glutamine, 1 mM sodium pyruvate and nonessential amino acids, 55 mM beta-mercaptoethanol, 20 mM HEPES, 100 U/mL penicillin, 100 μg/mL streptomycin, 30 μg/mL DNase I, and 0.5 mg/mL Liberase TL purified enzyme blend, Roche). Tissue isolated from mice treated with medroxyprogestrone acetate or low dose beta-estradiol were incubated for 25 minutes at 37^°^C and 5% CO_2_ and tissue isolated from mice treated with high dose beta-estradiol were incubated for 45 minutes at 37^°^C and 5% CO_2_. The digested tissue was then filtered through a 70 μM cell strainer, and the pellet was treated with ACK lysis buffer.

### In vitro stimulation of lymphocytes

To evaluate basal cytokine production, single-cell suspensions from uterine horns were cultured *ex vivo* for 2.5 hours at 37^°^C in 5% CO_2_ in complete medium (RPMI 1640 supplemented with 10% fetal bovine serum, 2 mM L-glutamine, 1 mM sodium pyruvate, 1 mM nonessential amino acids, 20 mM HEPES, 100 U/mL penicillin, 100 μg/ mL streptomycin, and 50 mM β-mercaptoethanol) containing 50 ng/mL of phorbol myristate acetate (PMA) (Sigma-Aldrich) and 5 μg/mL of ionomycin (Sigma-Aldrich), and a 1:1000 dilution of brefeldin A (GolgiPlug, BD Biosciences).

### Flow Cytometry

Single-cell suspensions were incubated for 30 minutes at 4^°^C in Hank’s buffered salt solution (HBSS) with combinations of fluorophore-conjugated antibodies against the following surface markers for a lymphoid panel: CD45 (clone - 30-F11, BD), NK1.1 (clone - PK136, BD), TCRβ (clone - H57-597, BD), CD90.2 (clone - 30-H12, BD), CD4 (clone - RM4/5, Biolegend), NKp46 (clone - 29A1.4, Biolegend), CD49a (clone - Ha31/8, BD), TCRγδ (clone - GL3, Invitrogen), CD8β (clone - H35-17.2, Invitrogen). Single-cell were incubated for 30 minutes at 4° in HBSS with combinations of fluorophore-conjugated antibodies against the following surface markers for a myeloid panel: CD45 (clone - 30-F11, BD), CD103 (clone - M290, BD), MHC-II (clone - 2G9, BD), CD11b (clone - M1/70, BD), CD64 (clone - X54-5/7.1, Biolegend), F4/80 (clone - BM8, Invitrogen), Ly6C (clone - HK1.4, Biolegend), CD90.2 (clone - 30-H12, BD), Ly6G (clone - 1A8, Biolegend), CD11c (clone - N418, Biolegend), B220 (clone - RA3-6B2, Biolegend).

To exclude dead cells, single cell suspensions were incubated with LIVE/DEAD Fixable Blue Dead Cell Stain Kit (Invitrogen Life Technologies) according to kit instructions. Cells were then fixed and permeabilized with the Foxp3/Transcription Factor Staining Buffer Set (eBioscience) and cells were incubated with intracellular antibodies at 4°C for 1 hour at room temperature. Combinations of fluorophore-conjugated antibodies against the following intracellular markers for a lymphoid panel include: Eomes (clone - Dan11mag, Invitrogen), Ki67 (clone - SolA15, Invitrogen), RORγT (clone - Q31-378, BD), T-bet (clone - 4B10, Biolegend), Foxp3 (clone - FJK-16s, Invitrogen), GATA3 (clone - TWAJ, Invitrogen), IL-17 (clone - TC11-18H10.1, Biolegend) and IFNγ (clone -XMG1.2, Invitrogen). Each staining was performed in the presence of purified anti-mouse CD16/32 (clone 93) and the extracellular staining was performed in the presence of BD Horizon™ Brilliant Stain Buffer. Cell acquisition was performed on a spectral flow cytometer (Cytek Aurora) using SpectroFlo software (Cytek v.1.1) and analyzed using FlowJo Software (BD).

### Single Cell RNA sequencing

Littermate WT (n=4) or IFNLR KO mice (n=3) were treated with medroxyprogesterone acetate, and the uteri were collected and digested as described above. Isolated cells were stained with antibodies against surface markers and TotalSeq-A hashtag oligonucleotide (HTO) antibodies (Biolegend) (**Table S1**). Hashtags with not enough reads (due to insufficient staining) were excluded from the downstream analysis. Cells were sorted and pooled together on a Sony MA900 sorter as live (DAPI-) EpCam^+^CD45^-^ or as live (DAPI-) and CD45^+^. The Chromium Single Cell Controller (10X Genomics) was overloaded with one lane containing ∼10,000 EpCam^+^ cells (∼1250 cells per sample) and another lane containing ∼22,000 CD45^+^ cells (∼2750 cells per sample) per lane. Libraries were prepared using a Chromium Single Cell 5’ Reagent kit following the manufacturer’s guidelines. Libraries were sequenced on an Illumina Nextseq2000 (Next Seq 1000/2000 P2 XLEAP-SBS 200 cycle kit, Illumina), with the EpCam^+^ library run on two flow cells and the CD45^+^ library run on four flow cells. Illumina files were converted to the FASTQ format using bcl-convert v4.2.7 command. 10x Genomics Cell Ranger v8.0.1 was used to process gene expression and hashtag oligonucleotide (HTO) data. First, the “cellranger mkref” command was used to generate a custom reference based on the mm10 (GRCm39) mouse genome primary assembly and the Ensembl release 113 protein-coding gene annotations. Next, “cellranger multi” was employed to generate gene expression and HTO count matrices.

The sequencing quality was high in the EpCam^+^ library, with > 95.6 % of bases in the barcode and UMI regions having a Q30 quality score and > 93% of bases in the RNA reads having a Q30 quality score. The median gene count was 2,441 with a sequencing saturation of 61% for the EpCam^+^ library and about 85.5 % of EpCam^+^ reads mapped confidently to the transcriptome. The mean read count per cell was ∼ 21,517 in the EpCam^+^ library. For the EpCam^+^ library only singlet cells with less than 5% mitochondrial contamination were used for downstream analysis, leaving 6,038 cells. The sequencing quality was high in the CD45^+^ library with > 95.6 % of bases in the barcode and UMI regions having a Q30 quality score and > 90.3% of bases in the RNA reads having a Q30 quality score. The median gene count per cell was 1,293 with a sequencing saturation of 86.13% for the CD45^+^ library and about 67.86% of CD45^+^ reads mapped confidently to the transcriptome. The mean read count per cell was ∼36,627 in the CD45^+^ library. For the CD45^+^ library only singlet cells with less than five percent mitochondrial contamination were used for downstream analysis, leaving 6,945 cells.

### scRNAseq data analysis

Downstream scRNAseq analyses were conducted using Seurat v5.1.0^111^. HTO demultiplexing was performed on centered log-ratio (CLR)-transformed HTO counts using the “HTODemux” function, with “positive.quantile = 0.99” and “nsamples = total number of cells”. Cells classified as “Singlet” in the “HTO_classification.global” output of “HTODemux” were retained for further analysis. Quality control filtering excluded cells with >5% mitochondrial gene expression, <500 detected genes, or >5000 detected genes. Remaining cells were normalized on a per-sample basis using the “SCTransform” function with default parameters. Dimensionality reduction was performed using “RunPCA”, followed by “RunUMAP” based on the top 30 principal components. Cell clustering was performed using “FindNeighbors” and “FindClusters”, with a resolution parameter of 0.3 for EPCAM^+^ cell sorts and 0.35 for CD45^+^ cell sorts. Cluster marker genes were identified using “FindAllMarkers” with the following parameters: “min.pct = 0.25”, “logfc.threshold = 0.5”, and “test.use = wilcox.” Cell type annotations of clusters were guided by the results of identified marker genes. Specifically, cluster identities were assigned by comparing these marker genes to known cell type specific markers as previously described and outlined in **Table S2**. Differential gene expression was assessed using the ‘FindMarkers’ function with ‘min.cells.group’ = 3, ‘min.pct’ = 0.01, ‘logfc.threshold’ = 0.1, and ‘test.use’ = ‘wilcox’. Genes were considered differentially expressed if they exhibited a Bonferroni-adjusted *p*-value < 0.05 and an absolute log2 fold change greater than 1. Gene Ontology (GO) enrichment analysis was performed on downregulated genes, defined as those with a log2 normalized fold change less than –2, using clusterProfiler v4.12.6 (https://doi.org/10.1016/j.xinn.2021.100141), while focusing on Biological Process ontology terms.

### Quantification and statistical analysis

Groups were compared with Prism V10.5 software (GraphPad) using the two-tailed unpaired Student’s t test, one-way analysis of variance (ANOVA) with Holm-Šidák multiple-comparison test, or two-way ANOVA with Holm-Šidák multiple-comparison test where appropriate. Where appropriate, outliers were identified and censored using the ROUT method. Differences were considered statistically significant when p ≤ 0.05.

## Funding

This research was supported by the Intramural Research Program of the National Institutes of Health (NIH). The contributions of the NIH authors are considered Works of the United States Government. The findings and conclusions presented in this paper are those of the authors and do not necessarily reflect the views of the NIH or the U.S. Department of Health and Human Services. ES, AR, EAG, CM, and IS are supported by the Division of Intramural Research of NIAID (NIAID; 1ZIA-AI001132, 1ZIA-AI001121, and 1ZIC-AI001233). ES is in part supported by the National Institute of General Medical Sciences Postdoctoral Research Associate Fellowship (1FI-2GM150424).

## Author Contributions

Conceptualization: ES, YB

Methodology: ES, YB, IS, EAG, JPC

Investigation: ES, AR, EAG, CM, JPC

Visualization: ES, YB, IS, AR, CM,

Supervision: ES, YB, IS

Writing - original draft: ES, YB, IS

Writing - review and editing: ES, YB, IS, AR, EAG, JPC, CM

## Competing Interests

The authors declare no competing interests.

## Data and materials availability

scRNAseq were deposited into the Gene Expression Omnibus (GEO) data repository (GSE314086). All data are available in the main text or the supplementary materials.

## Acknowledgments

We thank K. Beacht, E. Lewis, the NIAID animal facility, and the NIAID Microbiome Program’s gnotobiotic animal facility for technical support. Finally, we thank all the Belkaid and Sereti laboratory members for their generous support and providing feedback on this project. Some figures were created with BioRender.com.

---

**Figure S1.**
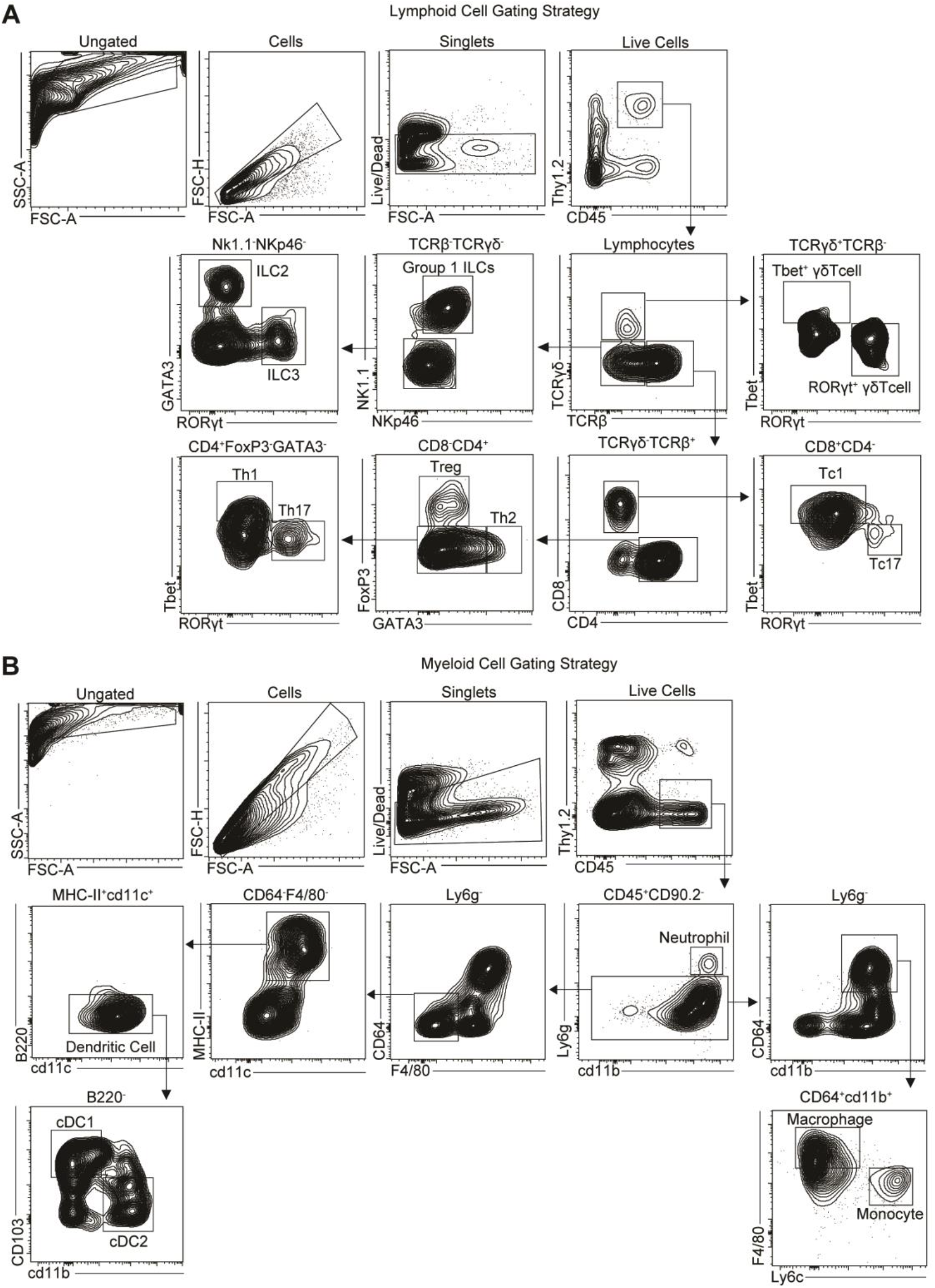
Gating Strategy. (A, B) Representative FACS plot showing (A) lymphoid and (B) myeloid gating strategy.

**Figure S2.**
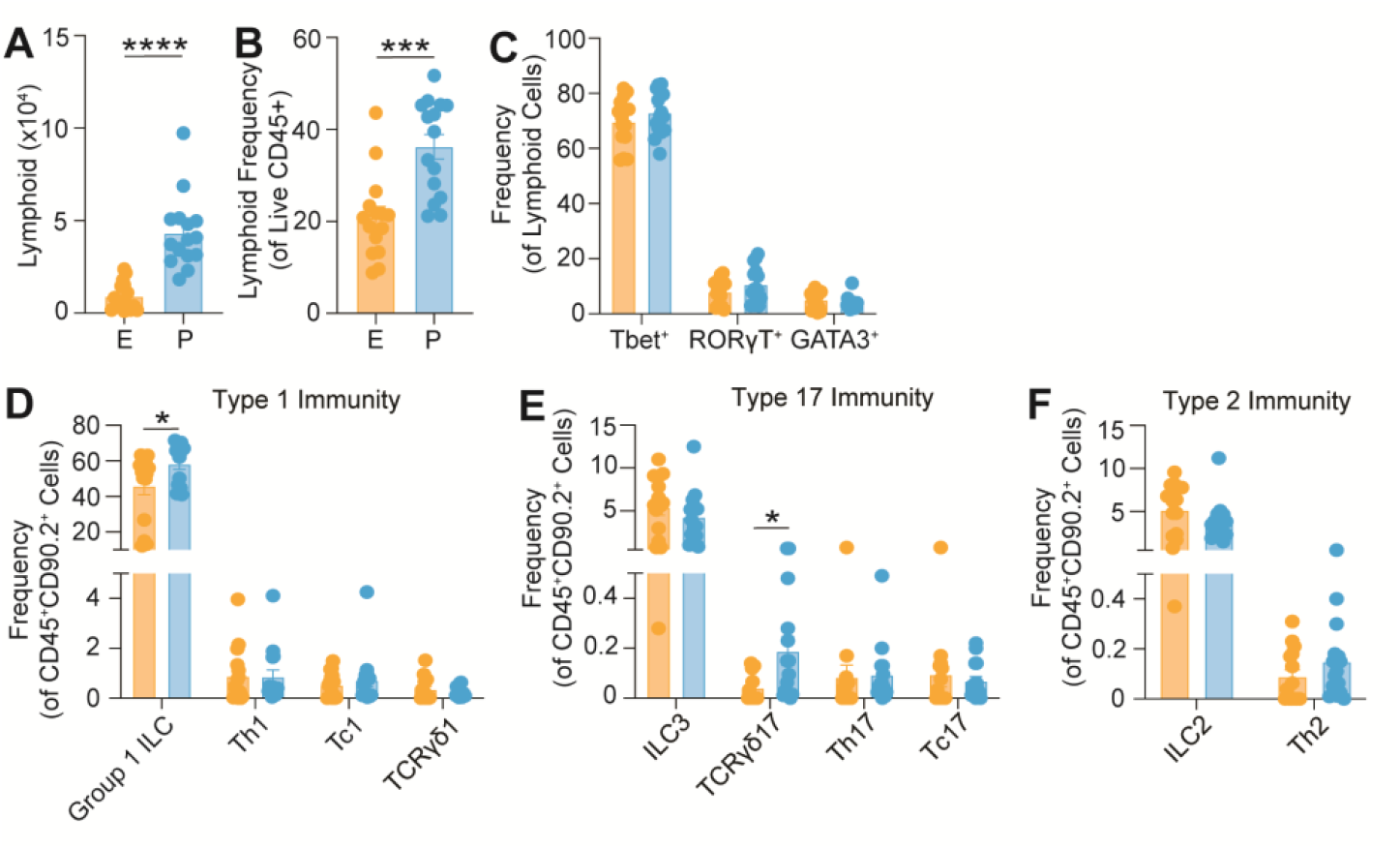
(associated with Figure 1). (A, B) Quantification of absolute cell number and cell frequency (of Live CD45^+^) of lymphoid cells from uteri of mice treated with estrogen (E, orange) or progesterone (P, blue). (C) Quantification of cell frequency (of CD45^+^CD90.2^+^) of Tbet^+^, RORγT^+^, or GATA3^+^ lymphoid cells from mice treated with estrogen or progesterone. (D, E, F) Quantification of cell frequency (of CD45^+^CD90.2^+^) of the indicated cell types. (A-F) Each dot represents an individual tissue with mean ± SEM from at least three biological replicates. Significance was determined by an unpaired Student’s t-test. * denote p-val<0.05 *** denote p-val<0.001 **** denote p-val<0.0001.

**Figure S3.**
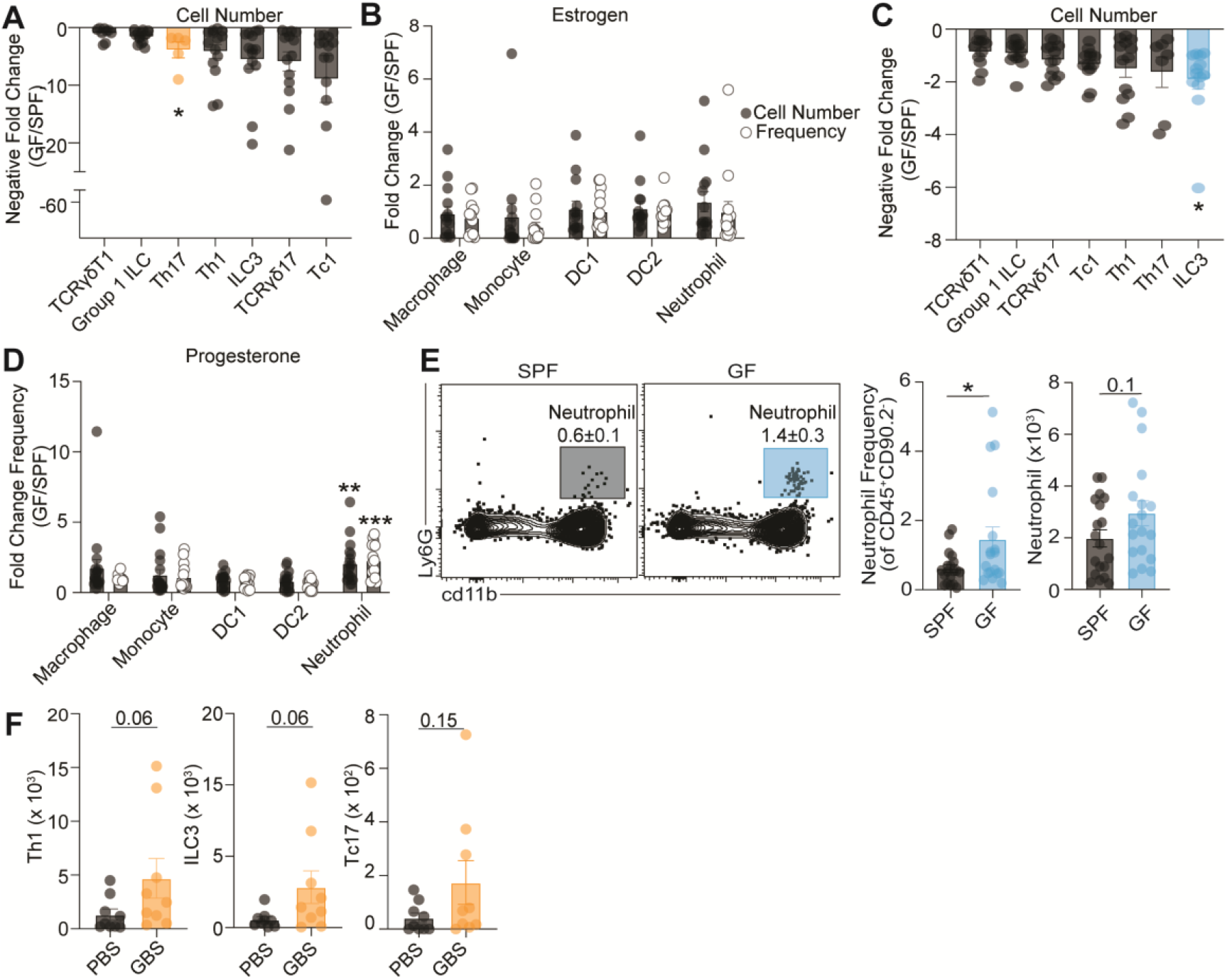
(associated with Figure 1). (A, C) Quantification of type 1 and type 17 immune cells as negative fold change in cell frequency (of CD45^+^CD90.2^+^ cells) in GF animals compared to SPF animals during (A) estrogen or (C) progesterone treatment. (B, D) Quantification of indicated myeloid cells as fold change in absolute cell number and cell frequency (of CD45^+^CD90.2^+^ cells) in GF animals compared to SPF animals during (B) estrogen or (D) progesterone treatment. (E) Left, representative FACS plots showing frequency of neutrophils in SPF and GF animals treated with progesterone. Right, quantification of absolute neutrophil cell number and frequency in SPF and GF animals treated with progesterone. (F) Quantification of absolute cell number of Th1, ILC3, and Tc17 cells. (A-D, F) Each dot represents an individual tissue from at least three biological replicates with mean with ± SEM. Significance determined by an unpaired Student’s t-test. * denote p-val<0.05 ** denote p-val<0.01*** denote p-val<0.001.

**Figure S4.**
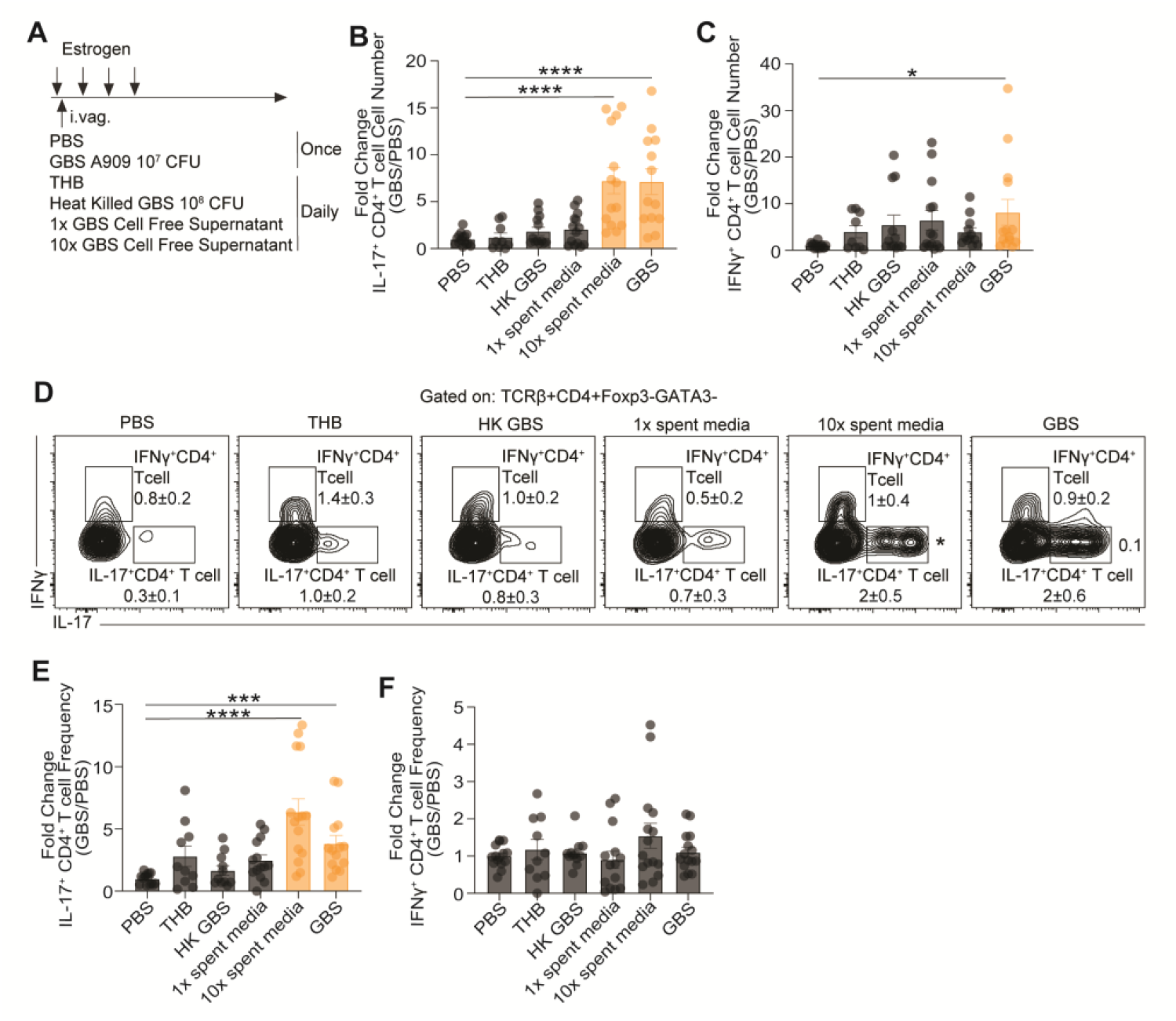
(associated with Figure 1). (A) Experimental schematic showing one time administration of live GBS and daily administration of heat killed GBS, 1x GBS spent media, 10x GBS spent media, and bacterial growth media (THB). (B, C) Quantification of (B) IL-17^+^ CD4+ T cell and (C) IFNγ^+^ CD4+ T cell numbers as fold change in absolute cell number in GBS treated animals relative to mock treated animals. (D) Representative FACS plots displaying frequency of uterine IFNγ^+^ or IL-17^+^ CD4^+^ T cells at 21 dpa with mock, GBS, heat killed GBS, 1x GBS spent media, and 10x GBS spent media. (E, F) Quantification of (E) IL-17^+^ CD4^+^ T cell and (F) IFNγ^+^ CD4^+^ T cell number as fold change in cell frequency (of CD4^+^ T cells) in GBS treated animals relative to mock treated animals. (B, C, E, F) Each dot represents an individual tissue from at least two biological replicates with mean with ± SEM. Significance determined by a one-way ANOVA with multiple comparisons relative to PBS treated mice. * denote p-val<0.05 *** denote p-val<0.001 **** denote p-val<0.0001.

**Figure S5.**
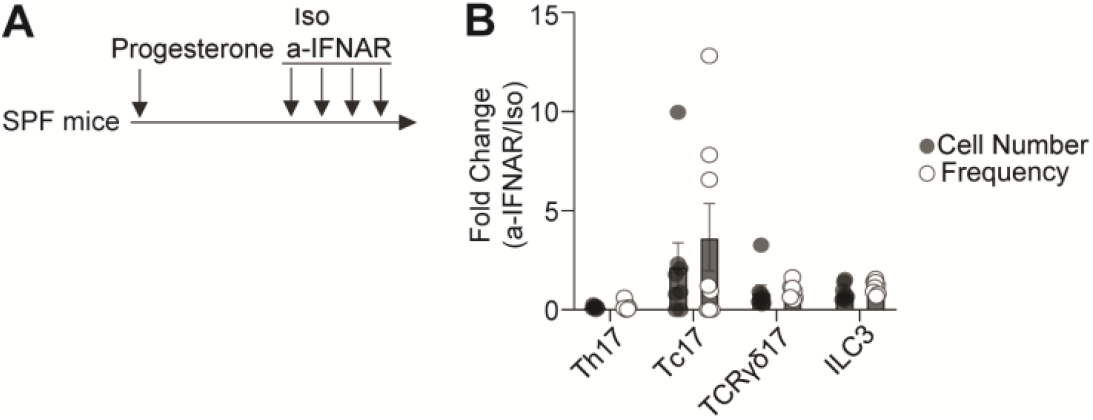
(Associated with Figure 2). (A) Experimental schematic depicting isotype control (Iso) or anti-IFNAR neutralizing antibody (a-IFNAR) treatment following progesterone treatment of SPF mice. (B) Quantification of indicated lymphoid cell types as fold change of absolute cell number and cell frequency (of CD45^+^CD90.2^+^ cells) in animals treated with a-IFNAR compared to Iso treated animals. Each dot represents an individual tissue from at least two biological replicates with mean ± SEM. (B) Significance determined by an unpaired Student’s t-test.

**Figure S6.**
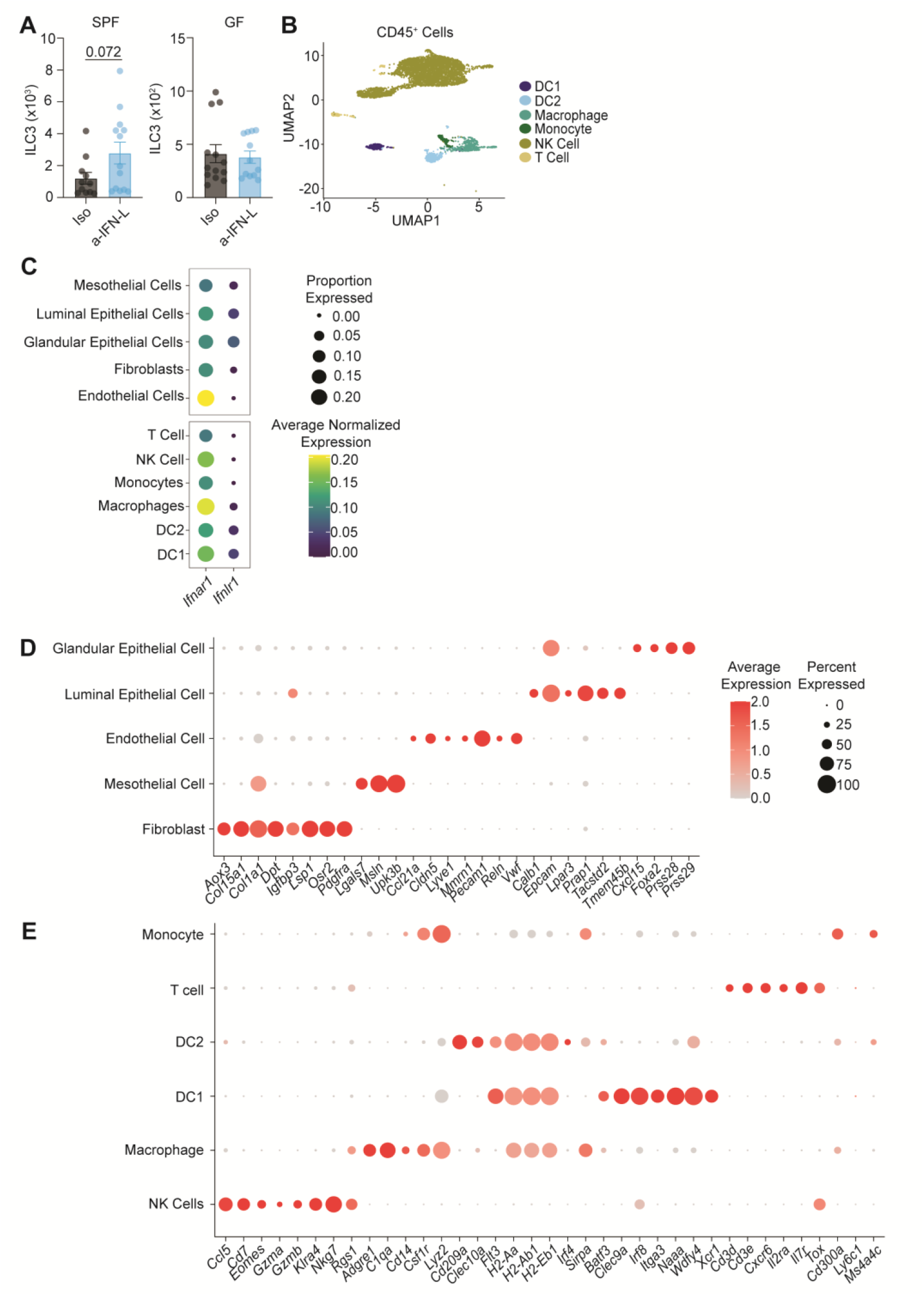
Associated with Figure 2. (A) Quantification of uterine ILC3 absolute cell number in SPF and GF animals treated with progesterone and Iso or a-IFN-L. Each dot represents an individual tissue from three biological replicates with mean ± SEM. Significance was determined by an unpaired Student’s t-test. (B) UMAP projection of CD45^+^ cells sorted from WT and IFNLR KO uteri. (C) Dot plot showing expression of *Ifnar1* or *Ifnlr* in WT cells from the EpCam^+^ and CD45^+^ library. (D, E) Dot plot showing marker gene expression for each cell type from the (D) EpCam^+^ or (E) CD45^+^ library.

**Table S1.**
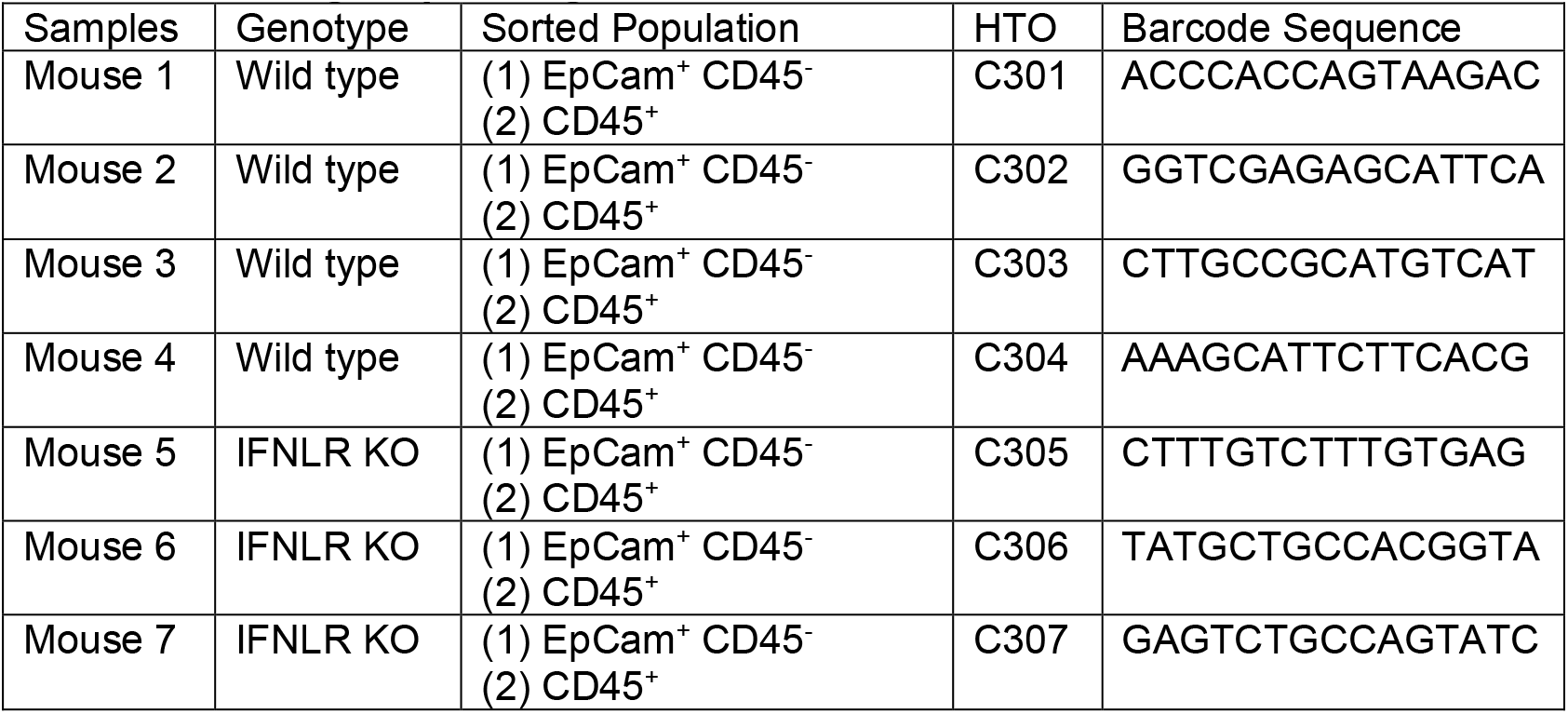
Hashtag Sequencing information.

**Table S2.** Cell type specific marker genes.

| **Cell type** | **Cell type abbreviation** | **Marker** | **Reference** |
| --- | --- | --- | --- |
| Glandular epithelial cell | GE | CXCL15, Foxa2, Prss28, Prss29 | 1, 2, 3 |
| Luminal epithelial cell | LE | Calb1, Tacstd2, EpCam, Lpar3, Prap1, Tmem45b | 2, 3, 4 |
| Mesothelial cell | M | Msln, Upk3b, Lgals7 | 5-7 |
| Fibroblast | F | Dpt, Col15a1, Lsp1, Igfbp3, Aox3, Col1a1, Pdgfra, Osr2 | 8, 9 |
| Endothelial cell | E | Pecam1, Cldn5, Vwf, Lyve1, Mmrn1, Ccl21a, Reln | 8-10 |
| Natural Killer cell | NK | Klra4, Rgs1, Cd7, Nkg7, Gzma, Gzmb, Ccl5, Eomes | 8, 9 |
| Macrophage | Mac | Csf1r, Cd14, C1qa, Adgre1, Lyz2, | 8, 9 ,11 |
| Dendritic cell type 2 | DC2 | H2-Ab1, H2-Eb1, H2-Aa, Cd209a, Clec10a, Sirpa, Irf4, Flt3 | 12 |
| Dendritic cell type 1 | DC1 | Clec9a, Wdfy4, Xcr1, Itgae, Naaa, Irf8, Flt3, Batf3 | 12 |
| T cell | T | Il7r, Cd3e, Cd3d, Il2ra, Cxcr6, Tox | 8, 9 |
| Monocyte | Mo | Ly6c, Cd300a, Adgre1, Lyz2, Ms4a4c | 9 ,11 |

## Notes

### Competing Interest Statement

The authors have declared no competing interest.

